# TALLSorts: a T-cell acute lymphoblastic leukaemia subtype classifier using RNA-seq expression data

**DOI:** 10.1101/2023.04.05.535648

**Authors:** Allen Gu, Breon Schmidt, Andrew Lonsdale, Lauren M. Brown, Teresa Sadras, Paul G. Ekert, Alicia Oshlack

## Abstract

T-cell acute lymphoblastic leukaemia (T-ALL) is an aggressive and heterogenous haematological malignancy affecting both children and adults. T-ALL subtype identification is an emerging area of active research, with several recent studies proposing potential subtypes based on transcriptomic and genomic analyses. Here we present TALLSorts, a machine-learning bioinformatic tool which classifies T-ALL samples by using bulk RNA sequencing (RNA-seq) data. Trained on four international cohorts totalling 264 samples, TALLSorts exhibits excellent accuracy when tested on holdout and independent test sets. TALLSorts is publicly available for use and will be constantly updated as the field of T-ALL classification further develops.

## Main text

T-cell acute lymphoblastic leukaemia (T-ALL) is a rare and aggressive haematological malignancy affecting both children and adults^1,2^. T-ALL subtype-identification is an emerging area of active research; as recently as 2016, the World Health Organisation (WHO) suggested only one provisional distinct T-ALL classification: the early T-cell precursor (ETP)^3^.

However, recent revisions by the International Consensus Classification in 2022 further subclassified ETP based on *BCL11B* deregulation, while introducing eight provisional classifications for non-ETP T-ALL^4,5^. These include subtypes characterised by rearrangements in *TAL1/2, TLX3, TLX3, LMO1/2, SPI, NKX2* and *BHLH*, and dysregulation of *HOXA*^4,5^. Other recent work by Dai *et al*., using gene expression analysis of eight international cohorts of adult and paediatric T-ALL samples, similarly developed classifications based on molecular abnormalities in *TAL1, TLX1, TLX3, NKX2, LMO1/2, LYL1* and *SPI*, while subclassifying *HOXA*-dysregulated samples by the presence of *KMT2A* or *MLLT10* fusions^6^, resulting in ten subtypes. Furthermore, Brady *et al*. recently described ten subtypes, defined again by *TAL1/2, TLX1, TLX3, NKX2, LMO1/2, SPI* and *HOXA* abnormalities, along with *BCL11B* abnormalities^7^ and an otherwise non-specified group^8^.

As with other leukaemias, understanding the heterogeneity within T-ALL holds clinical significance as T-ALL subtypes may influence risk stratification and treatment decisions. For instance, *TLX1* and *TLX3* lesions indicate good prognosis, TAL1/2 lesions indicate intermediate prognosis, and the rare *SPI* rearrangement subtype indicates very poor prognosis^5^.

Considering the recent expansion of T-ALL subtype classifications and the potential clinical utility, we developed TALLSorts, a machine-learning classifier algorithm which uses bulk RNA sequencing (RNA-seq) gene expression data to attribute subtypes to T-ALL samples. TALLSort’s architecture is similar to ALLSorts, an analogous B-ALL subtype classifier which we have previously published^9^. TALLSorts (in its current iteration) accepts an expression matrix of RNA-seq reads aligned to the hg19 reference genome. This data is then analysed by pre-trained logistic regression models employing the one-vs-rest strategy for each subtype. Each sample is then assigned a probability score for each subtype, with a positive identification made if one or several subtypes exceed the 50% threshold (see Supplementary Information for further details). Logistic regression models were built using the *scikit-learn* package in Python^10^.

In building a training set, we selected publicly-available RNA-seq expression datasets comprising 376 T-ALL samples from four international cohorts: 26 samples from the Royal Children’s Hospital in Australia^11^, 265 samples from the Therapeutically Applicable Research To Generate Effective Treatments (TARGET) program in the United States^12^, and two cohorts of 25 and 60 samples respectively from the Saint-Louis Hospital in France^13,14^.

On these four cohorts, we utilised ComBat^15^ and RUVseq^16^ to remove batch effects. We then performed dimension reduction and unsupervised clustering on gene expression using the Uniform Manifold Approximation and Projection (UMAP) and density-based spatial clustering (DBSCAN) methods respectively^17,18^, yielding eight distinct clusters (Figure 1A).

**Figure 1.**
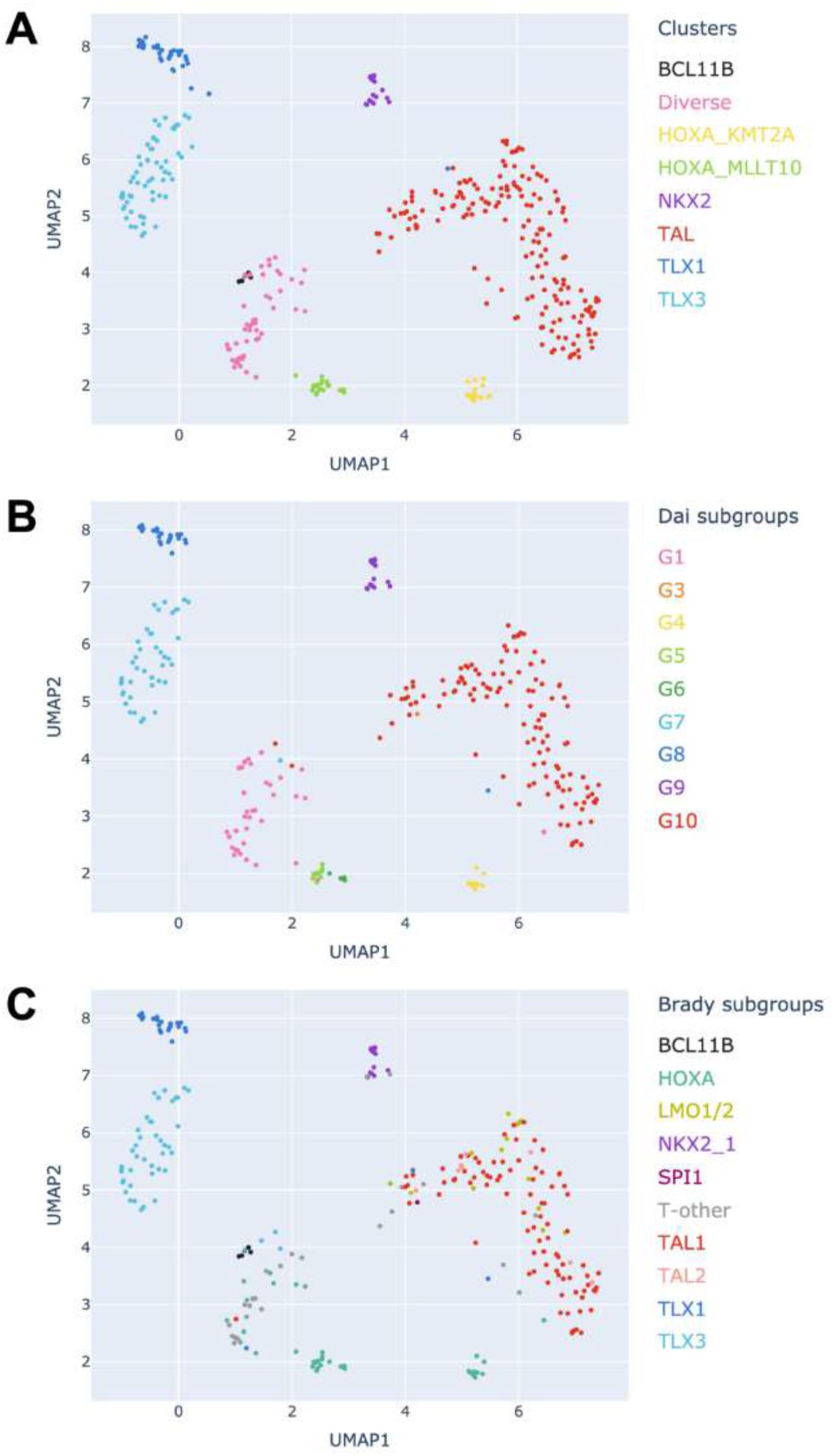
Identification of subtypes through clustering and dimension reduction. (A) UMAP dimension reduction and DBSCAN clustering of samples used for training and holdout testing (n=376), coloured by clusters with cluster labels informed by Dai *et al*. and Brady *et al*. (B-C) TARGET samples (n=265) within our clustering coloured by subtype labels assigned by Dai *et al*. (B) and Brady *et al*. (C).

As the TARGET cohort was analysed separately by Dai *et al*. and Brady *et al*., we compared their TARGET sample subtype labelling to our clusters, where we found an almost direct correspondence (Figure 1B-C). Considering each cluster as a distinct subtype, we assigned each of our discovered clusters with an appropriate subtype name reflecting the likely underlying pathology. This largely coincided with labels from Dai *et al*. and Brady *et al*. Specifically, our eight subtypes correspond to transcriptomic landscapes associated with *BCL11B* deregulation; *NKX2* overexpression; *TAL* deregulation; *TLX1* overexpression; *TLX3* overexpression; *HOXA* overexpression and fusions involving *KMT2A* or *MLLT10*; and a Diverse category. In the case of the *BCL11B* cluster, we manually separated it from the Diverse cluster by identifying samples within the *BCL11B* subgroup from Brady *et al*.’s classification system (Figure 1C). In the case of the Diverse subtype, its analogous groupings within both Dai *et al*. and Brady *et al*.’s systems suggest it is composed of samples with heterogeneous genetic profiles not otherwise captured by the other subtypes, and with lesions not otherwise defined.

The Rand index for concordance of TARGET samples between our cluster labels and Dai *et al*.’s subgroupings was 0.97, indicating a high degree of agreement. Similarly, the Rand index between our cluster labels and Brady *et al*.’s subgroupings was 0.87; this lower value was likely largely due to the absence of *LMO1/2* and the discrimination between *HOXA* samples within our labels, compared to that of Brady *et al*.

Validating TALLSorts on samples held out from training (n=112) demonstrated 97% accuracy (n=109) (Fig 2E). Samples classified into multiple subtypes were considered accurately labelled if the classified subtypes include the true label. Notably, of the 109 correctly labelled samples, all classified with greater than 70% probability, while 93 (85%) were correctly classified with a probability greater than 90%, thus demonstrating model confidence in its assignments.

**Figure 2.**
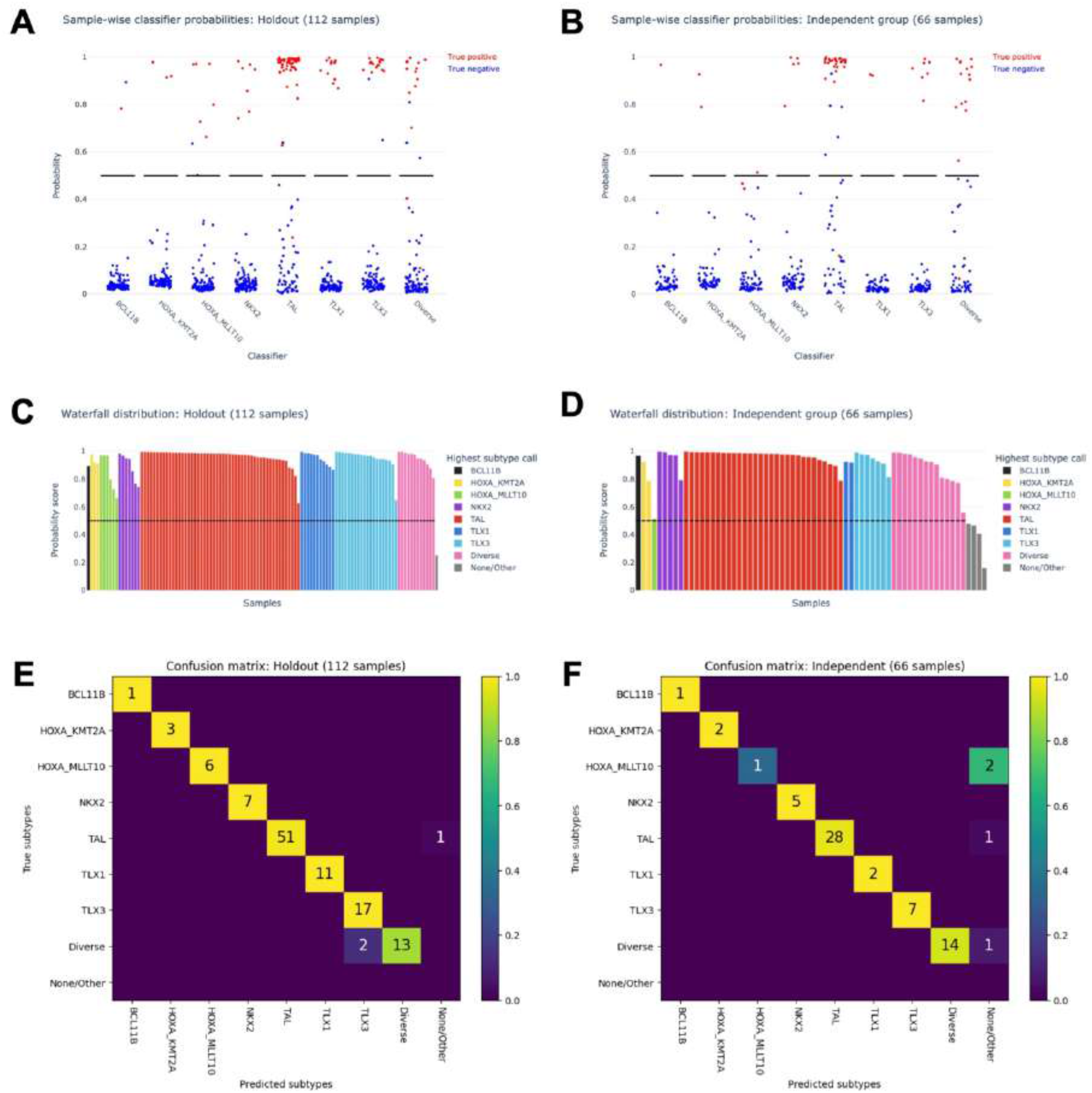
Validation of TALLSorts and example TALLSorts output figures. (A-B) Predicted probabilities for each sample (dot) by each subtype classifier (x-axis) within the holdout testing set (A) and the St. Jude/Ma-Spore independent testing set (B). A red dot indicates a sample labelled as the subtype being tested; a blue dot indicates a different label. The black line is the threshold of 50% probability; all samples above this line are classified as positive for the relevant subtypes. These probability plots are generated as output by the TALLSorts package, albeit without the true positive/negative colourings in the case of datasets without known truth values. (C-D) Waterfall plots for the holdout test set (C) and independent test set (D). Each vertical bar is a sample coloured by most likely subtype and with height proportional to its probability according to the classifier. Samples that do not exceed the 50% probability threshold for any subtype are labelled as “None/Other”. These waterfall plots are also generated as output by the TALLSorts package. (E-F) Confusion matrices for the holdout test set (E) and independent test set (F). The y-axes correspond to subtype ground truth, and the x-axes are the TALLSorts predicted subtypes. A sample predicted positive for multiple subtypes is considered correctly labelled if its true label is one of the predictions.

Next, we validated TALLSorts on 66 samples from two cohorts that were included in the study by Dai *et al*., but were incorporated into neither the training nor holdout data sets: a study from St Jude’s Children’s Research Hospital^19^ (n=41) and the Ma-Spore trial^20^ (n=25). Of the St Jude’s dataset, 30 of the 41 samples were also analysed in Brady *et al*. Truth values for these samples’ subtypes were determined by cross-referencing their Dai *et al*. and/or Brady *et al*. labels (Supp Figure S1). This dataset similarly exhibited good accuracy at 94% (n=62) (Fig 2F). Of these, 58 correctly-classified samples (94%) had a probability of greater than 70%, while 50 (81%) received greater than 90% probability. While these samples are independent from the training and holdout sets, we acknowledge the limitation that their truth values were informed by the labelling from Dai *et al*., which were also used to generate truth values for the training and holdout sets.

As mentioned previously, the multiclass logistic regression model of TALLsorts uniquely permits samples to be classified under multiple subtypes. This feature may prove beneficial in samples with more than one driver lesion, which may be only partially classified using decision-tree,or clustering-based algorithms as suggested by Brady *et al*., or Dai *et al*. respectively. Importantly, in the holdout dataset, we identified eight samples classified into multiple subtypes, and three samples similarly identified in the independent dataset.

The TALLSorts software will undergo active maintenance, including retraining, to ensure its subtype classifications remain current with the latest developments in T-ALL classification. Specifically, we expect that as more molecular studies investigate specific lesions and their underlying mechanisms and clinical outcomes, the subtype definitions will become further refined. At this stage we do not know the defining features of the Diverse classification which we expect contains several driving mechanisms. Similarly, as more samples are made available, more refined subtypes will be able to be discriminated from the gene expression data.

In conclusion, TALLSorts is an accurate, publicly-available algorithm that uses RNA-seq expression to classify T-ALL subtypes, where these subtypes reflect the latest research in T-ALL subclassification.

Further details regarding the construction and testing of TALLSorts is available in the accompanying Supplementary Information document. TALLSorts and its source code are available for download at https://github.com/Oshlack/TALLSorts.

## Acknowledgements

RCH tumour samples and coded data were supplied by the Children’s Cancer Centre Biobank at the Murdoch Children’s Research Institute and The Royal Children’s Hospital (https://www.mcri.edu.au/research/projects/childrens-cancer-centre-biobank). Establishment and running of the Children’s Cancer Centre is made possible through generous support by Cancer In Kids @ RCH (https://www.cika.org.au), The Royal Children’s Hospital Foundation and the Murdoch Children’s Research Institute. The authors acknowledge the support of the SCOR Grant (7015-18) from the Lymphoma and Leukemia Society and of Perpetual Trustees and the Samuel Nissen Foundation. AO is funded by an NHMRC Investigator Grant GNT1196256.

## Authorship contributions

A.G., B.S. and A.O conceptualised the study and created the methodology. P.G.E., L.M.B. and T.S. provided clinical and biological expertise. A.G. and B.S. wrote the software. A.G. and A.L. performed data extraction and sequence alignment. A.G. performed the formal analysis, and provided the original draft of the manuscript. A.G. and A.O. edited and reviewed the manuscript. All authors read and approved the manuscript.

