## Supplementary Information for "TALLSorts: a T-cell acute lymphoblastic leukaemia subtype classifier using RNA-seq expression data"

### Constructing the TALLSorts classifier

#### Training and testing data sources

To build the testing/holdout datasets, we used four T-ALL cohorts with publicly-available RNA-seq datasets.

| Cohort | Number of samples | Source |
|---|---|---|
| TARGET | 265 | Liu et al. (2017)[1] |
| GSE110633 | 60 | Verboom et al. (2018)[2] |
| GSE110677 | 25 | Buratin et al. (2020)[3] |
| Royal Children's Hospital (RCH) | 26 | Brown et al. (2020)[4] |

**Supplemental Table S1: Data sources used to build the classifier.**

The TARGET dataset is hosted by dbGaP under accession phs000218. The GSE110633 and GSE110677 datasets are hosted by NCBI GEO under the respective accessions. The RCH dataset is available at the European Genome-Phenome Archive under accession EGAS00001004212. The gene expression counts matrix for these samples were generated by STAR using the hg19 (GRCh37) reference genome.

#### Batch-effect removal

Prior to a clustering analysis, batch-effect removal was required to integrate samples from the four cohorts. Batch-effect removal was performed in two stages.

In the first-pass batch-effect removal process, we commenced with the raw counts matrix of training samples (376 samples and 51171 expressed RNA sequences including coding and noncoding genes) from the four training cohorts. However, before batch-effect removal, we filtered out the most cohort-correlated genes identified using a differential gene expression (DGE) analysis with the *edgeR* R package[5], which compared samples from each cohort against the others; for each of the four one-vs-rest cohort-wise comparisons, we selected the 1000 genes most correlated with cohort, resulting in 2857 genes (accounting for duplicates) to be filtered out. Batch-effect removal was next performed using the ComBat function of the R package *SVA*[6,7].

A second-pass batch-effect removal process was then performed. Firstly, a similar DGE analysis as the above was performed using *edgeR* to identify cohort-correlated genes; for each of the four one-vs-rest cohort-wise comparisons, we identified the 500 genes most correlated with cohort, which resulted in 1495 genes in total. Another DGE analysis was performed which similarly identified the genes most correlated with the preliminary subtypes identified post-ComBat analysis: for each of the eight one-vs-rest subtype-wise analyses, we amalgamated the 3000 most correlated genes, resulting in 14542 genes. We next removed all genes from the 1495 cohort-correlated genes list that were also in the 14542 subtype-correlated gene list. This gave a final list of 1279 genes that were correlated with cohort but not with subtype. We now performed batch removal using *RUVseq*[8], where the aforementioned list of 1279 genes served as a negative control against which the other genes were normalised.

The final result of the batch-effect removal process was a normalised gene counts matrix, which was used to identify and label subtypes through clustering. Critically, this normalised matrix was *not* used as input data in either training nor testing.

Code used for batch-removal is available upon request to the corresponding author.

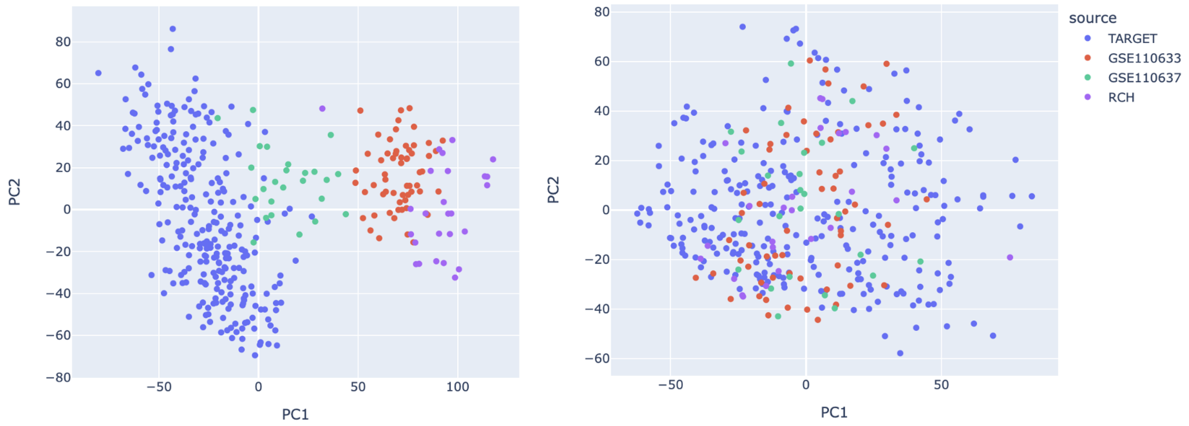


**Supplemental Figure S1: Batch effect removal results.** Principal component analysis (PCA) of gene expression counts for samples in the four training cohorts before (left) and after (right) batch-effect removal, demonstrating satisfactory integration of the cohorts.

### Identifying and labelling clusters

Beginning with the *RUVseq*-normalised counts matrix, we performed dimension reduction using the Uniform Manifold Approximation and Projection (UMAP)[9] method through the *umap* Python package. The data was projected to a three-dimensional space with parameters of *n_neighbours=10* and *min_dist=0*, with all other parameters at default values.

On this projection, the Density-Based Spatial Clustering of Applications with Noise (DBScan) algorithm[10] through the *scikit-learn* Python package[11] was performed with parameter *eps=0.7* and all others at default. This identified seven clusters (see Fig 1A). We compared how the TARGET samples within our clustering were classified by Dai *et al.*[12] and Brady *et al.*[13], which allowed us to create appropriate names for the cluster/subtypes. We were able to immediately name seven of the eight subtypes included in TALLSorts: *NKX2* overexpression*; TAL* deregulation*; TLX1* overexpression; *TLX3* overexpression; *HOXA* overexpression and fusions involving *KMT2A* or *MLLT10*; and a diverse category.

Notably however, comparison with the Brady *et al*. labelling demonstrated a group of five TARGET samples nested within the diverse category, which were identified as samples with *BCL11B* deregulation. We therefore manually labelled these samples as the *BCL11B* subtype, which was the eighth subtype included in TALLSorts.

| **TALLSorts subtype** | **Dai *et al.* labelling** | **Brady *et al.* labelling** |
|---|---|---|
| BCL11B | - | BCL11B |
| HOXA (KMT2A) | G4 | HOXA |
| HOXA (MLLT10) | G5; G6 | HOXA |
| NKX2 | G9 | NKX2_1 |
| TAL | G10 | TAL1; TAL2 |
| TLX1 | G8 | TLX1 |
| TLX3 | G7 | TLX3 |
| Diverse | G1 | T-other |

**Supplemental Table S2: TALLSorts subtype and the corresponding labels from Dai *et al.* and Brady *et al.***

Within Dai *et al.*'s groupings, G2 and G3 were not represented within TALLSorts; the former likely because G2 were composed of primarily adult samples while TALLSorts was primarily trained on paediatric samples, while G3 represented *SPI1*-fusions which which were not captured within our training set.

Within Brady *et al.*'s groupings, the *LMO1/2* label was not represented, as Fig 1C demonstrates the degree to which samples labelled as *LMO1/2* alterations were dispersed amongst *TAL1/2*-dysregulated samples; this raises the possibility that the *TAL* TALLSorts subtype may encompass samples with either *TAL1/2* and/or *LMO1/2* abnormalities. Furthermore, the *SPI1* label was also not represented.

The Rand index was used to examine the concordance between TALLSorts subtypes and each of the Dai et al. and Brady et al. groupings for the 265 TARGET samples, which were 0.97 and 0.87 respectively.

### Developing the Logistic Regression classifier

From each of the four labelled training cohorts (total n=376), 70% of the samples were randomly chosen to serve as the training set (n=264). The remaining 30% from each cohort were pooled into the holdout testing set (n=112).

### Filtering

Beginning with the un-normalised training counts matrix which contained 264 samples across 52,683 RNA-seq sequences, all Y-chromosome genes and the *XIST* gene were firstly removed. We then removed all transcripts that were not protein-coding genes or were mitochondrial genes according to Ensembl.

To filter low-expression genes, a gene was retained only if its count was greater than 5 in at least as many samples as there are samples in the lowest-populated subtype, which equalled 4 samples in the BCL11B subtype within the training set. This way, 16,147 genes were retained for training.

### Normalisation and scaling

Counts were converted to counts-per-million (CPM) values and normalised using the trimmed mean of M-values (TMM) method[14]. The natural log was taken to generate log-CPM values. Gene-wise standardisation was performed such that all genes' log-CPM values had zero mean and unit standard deviation.

### Logistic regression

For each of the eight subtypes, a logistic regression model was trained using the *LogisticRegression* class within the *scikit-learn* Python package[11], with parameters of tol=0.0001, max_iter=10000, the SAGA solver. Regularisation was achieved with L1 penalty and the value C=0.2, which was a relative measure of regularisation strength. Default options were used otherwise.

The effect of regularisation was to help prevent overfitting by removing as many uninstructive genes as possible. The following table shows the number of genes with non-zero coefficients for the subtype classifier in question.

| **Subtype** | **Number of genes post-regularisation** |
|---|---|
| BCL11B | 23 |
| HOXA_KMT2A | 36 |
| HOXA_MLLT10 | 53 |
| NKX2 | 20 |

| | |
|---|---|
| TAL | 64 |
| TLX1 | 31 |
| TLX3 | 26 |
| Diverse | 59 |

**Supplemental Table S3: Number of genes with non-zero coefficients post-regularisation for each subtype classifier.**

A full list of classifier coefficients and intercepts for each logistic regression model may be found in Supplemental Table S6. Specifically, the probability of a positive classification was given by

$$p = \frac{1}{1 + \exp(-\beta_0 - \sum_i \beta_i x_i)}$$

where $x_i$ was the standardised, normalised log-CPM count of the $i$-th gene; $\beta_i$ was the fitted coefficient for the $i$-th gene; and $\beta_0$ was the fitted model intercept term[11].

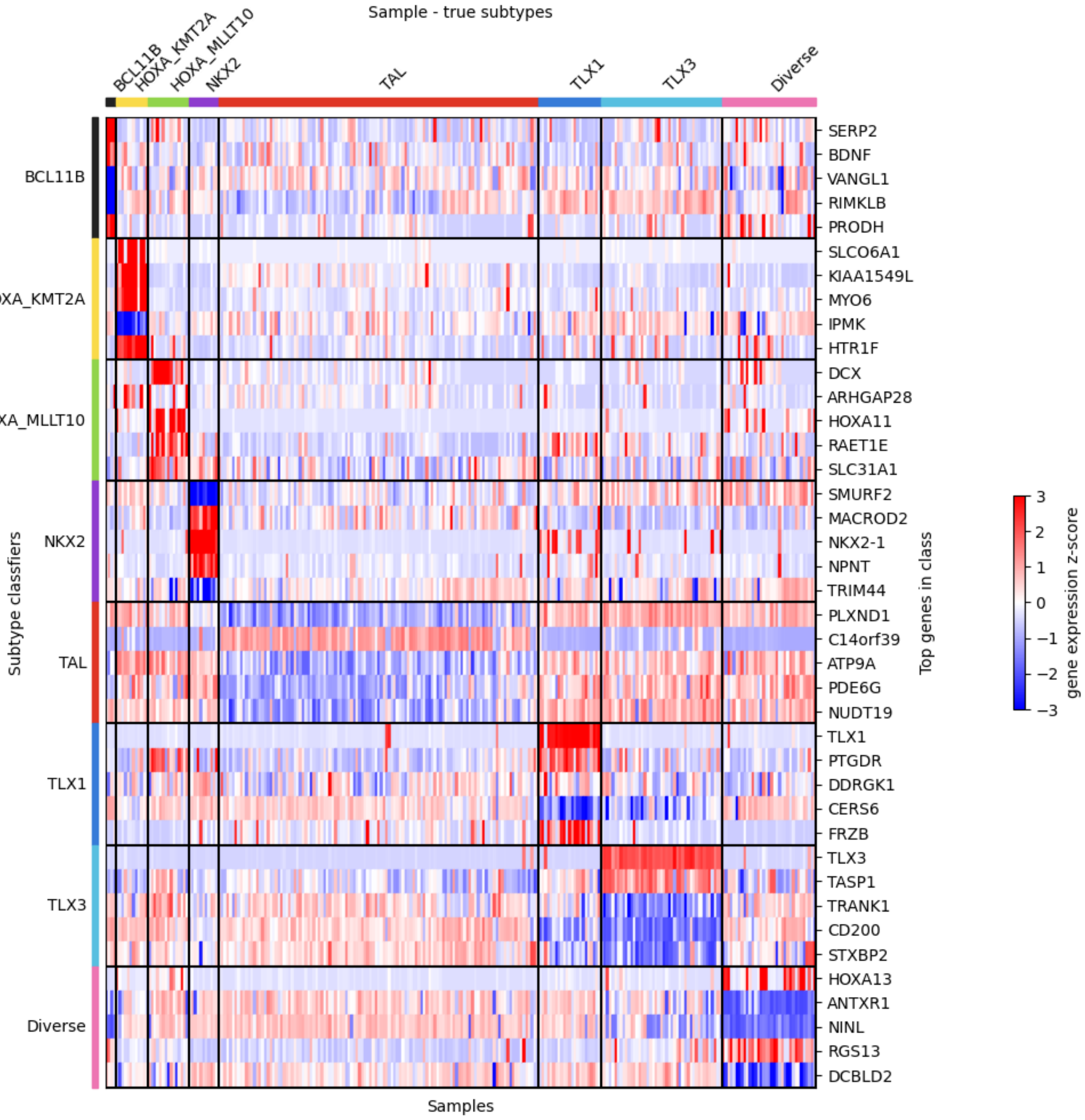


**Supplemental Figure S2: Heatmap demonstrating the most highly-correlated genes for each subtype.** Each small column represents a distinct sample within the training set (n=264). Samples are grouped by their "true" subtype as determined by the clustering algorithm.

### Validating and predicting with the TALLSorts classifier

#### Test sets

For this research letter, we employed two test sets. The first was the holdout set separated from the training set, containing the 112 samples that represented 30% of the total samples within the four training cohorts. The second was a pooled group of 66 samples from two international cohorts that were also analysed by Dai *et al*.

| Cohort | Number of samples | Source |
|---|---|---|
| MaSpore trial | 25 | Qian et al. (2017)[15] |
| St Jude's Children's Research Hospital | 41 | Autry et al. (2020)[16] |

**Supplemental Table S4: Datasets used as an independent test set.**

The MaSpore dataset is available at the European Genome-Phenome Archive under accession EGAD00001002151. The St Jude's dataset is available at NCBI GEO under accessions GSE115525 and GSE124824. The gene expression counts matrix for these samples were generated by STAR using the hg19 (GRCh37) reference genome.

#### Normalisation and standardisation

Similar to the training set, test sets or prediction sets were firstly converted to log-CPM and normalised using TMM. The normalised counts were then standardised; for a given gene count for a given sample, it was transformed by subtracting the mean count of that gene in the training set, then dividing the result by the standard deviation of that gene in the training set.

Genes in the test set that were not in the training set were removed from consideration in the classification step. Conversely, genes in the training set but not in the test set were appended to the test set with values of zero.

#### Prediction

The standardised counts matrix for a test set were then analysed by the pre-trained logistic regression models, each of which generated probabilities of the samples belonging to the respective subtypes. A threshold of 50% probability was set to distinguish positive and negative identifications. In the event of multiple subtypes being called, a sample was deemed to be correctly classified if its true subtype was among the called subtypes.
